# Why panmictic bacteria are rare

**DOI:** 10.1101/385336

**Authors:** Chao Yang, Yujun Cui, Xavier Didelot, Ruifu Yang, Daniel Falush

**Author notes:** Corresponding authors: D. F. These authors contributed equally to the article (Chao Yang, Yujun Cui).

## Abstract

**Background:** Bacteria typically have more structured populations than higher eukaryotes, but this difference is surprising given high recombination rates, enormous population sizes and effective geographical dispersal in many bacterial species.

**Results:** We estimated the recombination scaled effective population size *N*_*e*_*r* in 21 bacterial species and find that it does not correlate with synonymous nucleotide diversity as would be expected under neutral models of evolution. Only two species have estimates substantially over 100, consistent with approximate panmixia, namely *Helicobacter pylori* and *Vibrio parahaemolyticus*. Both species are far from demographic equilibrium, with diversity predicted to increase more than 30 fold in *V. parahaemolyticus* if the current value of *N*_*e*_*r* were maintained, to values much higher than found in any species. We propose that panmixia is unstable in bacteria, and that persistent environmental species are likely to evolve barriers to genetic exchange, which act to prevent a continuous increase in diversity by enhancing genetic drift.

**Conclusions:** Our results highlight the dynamic nature of bacterial population structures and imply that overall diversity levels found within a species are poor indicators of its size.

## Background

Bacteria are paragons of adaptability and make up more biomass than all organisms other than plants combined [1]. Many bacterial species have enormous population sizes, disperse effectively around the globe [2-5] and exhibit high rates of within-species homologous recombination [6]. Recombination progressively breaks down non-random associations between markers (linkage disequilibrium), so that in large populations, where pairs of individuals distantly related, linkage equilibrium is expected between all pairs of genomic sites. However, in most species for which data is available, there is substantial genome-wide linkage disequilibrium, indicating structuring of variation [7]. Here we propose a resolution of this paradox, namely that large bacterial populations accumulate diversity progressively until that diversity acts as an effective barrier to genetic exchange between lineages.

We examine the population structure of 21 bacterial species and find that only the Asian population of *Vibrio parahaemolyticus* is close to genome-wide linkage equilibrium. We find that this population has undergone a recent expansion. If the current population size was maintained over evolutionary timescales, it would lead to a greater than 30 fold increase in diversity, to levels higher than found in any well characterized bacterial species. We propose that other environmental species with large census population sizes, for example *Vibrio cholerae* or *Klebsiella pneumoniae*, may have been through a similar stage before accumulating diversity and that this diversity acted to generate barriers to recombination, either directly or via selective pressure to reduce recombination rates between genetically divergent lineages.

Specifically, we use pairwise genetic distances between strains calculated from core genome sequences to estimate the recombination scaled effective population size parameter *N*_*e*_*r*, where *N*_*e*_ is the effective population size and *r* is the recombination rate per site per generation, an approach briefly introduced by Cui et al [8]. Here, we test the method using simulated genomic datasets and find that it provides accurate estimates in constant size populations for values of the scaled recombination rate *R* (see below) of 5 or more, and responds to changes in population size or structure far more quickly than estimates of *N*_*e*_ based on nucleotide diversity. We apply the method to real genomic datasets from 21 species and find that some common species, such as *Escherichia coli*, have strikingly low estimates of *N*_*e*_*r*, implying that recombination is inefficient in distributing diversity across the species, while for others the fit of model to data is poor, highlighting deviations from the assumption of a freely recombining population. Only datasets from *V. parahaemolyticus* and *Helicobacter pylori* give *N*_*e*_*r* estimates greater than 100. The former has had a recent increase in population size, whereas the latter experienced repeated bottlenecks associated with geographical spread of its human host, implying that neither population is close to demographic equilibrium.

### Inference approach

Population genetics is a century-old discipline that provides a powerful set of theoretical and statistical inference tools with which to interpret patterns of genetic variation between closely related organisms. Central to the theory is the concept of a population, which in outbreeding eukaryotes is a set of organisms that share a common gene pool. The simplest models assume panmixia which means random mating of all individuals within a single closed population [9]. In most animals and plants mating is structured by geography, however even low levels of migration between locations, of the order of one individual per generation, can prevent differences from accumulating between local populations, making random mating a reasonable first approximation for many outbreeding species. Under neutral theory [10], the expected level of genetic diversity *π* depends on the per generation mutation rate *μ* and the effective population size *N*_*e*_, which is the number of individuals contributing to the gene pool in each generation [11].

Outbreeding eukaryotes receive genetic material from the gene pool when they are born and contribute to it when they reproduce. Individuals are more similar to immediate relatives than they are to other members of the population, but the proportion of the genome shared by descent from particular ancestors decays rapidly with each successive generation, declining from 1/2 for full siblings to 1/8 for first cousins and 1/32 for second cousins. In small random samples from populations with more than a few hundred individuals, shared recent ancestry is rare and can be neglected for many types of analysis.

Bacteria reproduce by binary fission and only receive material from the gene pool or contribute to it via homologous or non-homologous recombination [12]. Many cell divisions can take place between consecutive recombination events, and typically recombination only affects a small fragment of the genome. These properties mean that the concept of a gene pool or a population is less straightforward to define than for outbreeding organisms. It is still possible to estimate the effective population size *N*_*e*_ from the average nucleotide diversity *π*, based on the standard assumptions of population genetic theory, but the assumptions are both less reasonable and harder to test for bacteria than for outbreeding eukaryotes, in which population boundaries can be delineated empirically using well established methods [13]. For example, nucleotide diversity in a species like *E. coli* can be estimated for clonal lineages, phylogroups, species or for the *Escherichia* genus as a whole. These choices lead to very different values for *π* and hence for *N*_*e*_ and it is not obvious a priori which is most meaningful. All methods lead to estimates of effective population size that are many orders of magnitude smaller than the census number of bacteria [14].

Here, we take a different approach which is to estimate a scaled version of the effective population size, *N*_*e*_*r*, where *r* is the per generation rate at which a given site recombines. Note in particular that this parameter *r* is the product of the per initiation site recombination rate *ρ* and mean tract length *δ* used in many bacterial recombination models [15, 16]. High values of *N*_*e*_*r* indicate that the population structure is similar to that of outbreeding eukaryotes. Informally, in eukaryotic population each individual is the product of a separate meiosis and therefore genetically distinct. *N*_*e*_ is the number of genetically distinct individuals that contribute in each generation, which is typically in the thousands or millions. *N*_*e*_*r* is designed to measure an analogous quantity in bacteria, namely the number of genetically distinct organisms that contribute to the future bacterial gene pool. To this end, time is rescaled; *N*_*e*_ measures the rate of genetic drift per bacterial generation, while *N*_*e*_*r* measures the genetic drift in proportion to the time it takes for strains to become distinct from their ancestors by importing DNA by homologous recombination.

The recombination rate of *V. parahaemolyticus* is *r* =1.7 × 10^−4^ per site per year [17]. After *T* years of evolution, the expected proportion of recombined genome is e^-*rT*^, so that it takes around 4,000 years on average for half of the genome to recombine. Half of this time is approximately equivalent to a eukaryotic generation in the sense that two strains that shared a common ancestor 2,000 years ago will be about as related as siblings. This represents a very different time scale from that assumed based on bacterial generations. For example, *V. parahaemolyticus* is capable of replicating in less than ten minutes in appropriate conditions [18].

We estimate *N*_*e*_*r* using the pairwise genetic distances between strains, based on a number of simplifying assumptions which are likely to hold true in freely recombining bacterial populations but may break down in species where recombination rates are low or genetic exchange is structured by geography or lineage. Specifically, the calculations assume that recombination introduces many more substitutions than mutation, happens at the same rate throughout the genome and that each recombination event introduces unrelated DNA from the population into the imported stretch. If unrelated strains differ on average at *d*_unrelated_ nucleotides throughout the alignment, then the expected number of SNPs distinguishing strains with a common ancestor at time *T* in the past is *d* = *d*_unrelated_ (1 - e ^*-2rT*^).

To use these times to estimate the effective population size, we assume that the genealogy of clonal relationships is generated by a coalescent model with a constant population size *N*_*e*_ [19, 20]. This model generates expectations for the times in the past at which common ancestors of strains in a sample existed. Specifically, for a sample of *n* strains, there are *n* - 1 coalescent nodes. The age of the most recent node corresponds to the common ancestor of the two most closely related strains in the sample, while the (*n* - 1)^th^ node corresponds to the common ancestor of all the strains in the sample.

Coalescent theory implies that the expected time in the past at which the *m*^th^ most ancient coalescent event occurs is 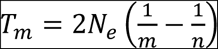. These times can be converted into expected genetic distances *d*_*m*_ using the formula in the previous paragraph. We use the UPGMA algorithm to obtain *n* - 1 coalescent distances from the *n*(*n* - 1)/2 pairwise genetic distances between strains and find the values of *N*_*e*_*r* and 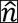 (effective sample size, see below) that gives the best fit between observed and expected distances for the *n* - 1 coalescent nodes.

Note that for bacteria with high recombination rate, the genome is likely to have been scrambled up sufficiently that strains will have inherited little or no DNA by direct descent from the common ancestor of the entire sample. This means that the genetic distances expected for the oldest coalescent events plateau at *d*_unrelated_. In graphical representations, it is convenient to show the coalescent events in chronological order with the oldest first, to aid comparisons between datasets with different sample sizes.

The model assumes that the strains are randomly sampled from a homogeneous population at a single time point, but pathogenic clones, epidemic outbreaks or strains from specific locations are often overrepresented in real data, leading to oversampling of very closely related isolates, relative to their frequency in the global population [7, 21]. Therefore, in addition to *N*_*e*_*r* we estimate a second parameter 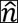 called the effective sample size, which is an estimate of the number of strains remaining when over-sampled clonally related strains are removed. For simulated data where sampling is random, 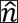 is correctly estimated to be very close to number of strains in the sample, so this additional parameter makes little difference to the inference. For real data, estimating this additional parameter often improves the qualitative model fit considerably.

## Results

### *N*_*e*_*r* can be estimated accurately for simulated data

Fig. 1 illustrates the effect of varying the recombination rate in genomes simulated using FastSimBac [22]. The simulations include 200 genomes of length 2 Mb under a coalescent model with constant effective population size *N*_*e*_. We fix the recombination tract length *δ* = 1,000 and vary the rate of initiation of recombination events *ρ* per 2*N*_*e*_ generations (0.001 to 0.1), so that the scaled recombination rate 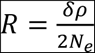 varies by two orders of magnitude, from 0.5 to 50. We also fix the rate of initiation of recombination event *ρ* (0.01) and vary the recombination tract length *δ* (100 to 10,000). Both methods give similar results, for low scaled recombination rates, the phylogenetic tree is highly structured but becomes progressively bushier as *R* increases, and gives the impression of being a nearly perfect star for *R* = 50. For high *R*, the estimated *N*_*e*_*r* is close to *R* and the observed genetic distances are well-fit by the model. For lower *R*, the fit is less good and the estimate of *R* provided by *N*_*e*_*r* is also less accurate, although it remains of the right order. For these parameter values, the size and shape of the phylogenetic tree are highly variable between runs, and the poorer fit reflects this stochasticity as well as greater inaccuracy in the approximations made in converting between genetic distances and coalescent times.

**Figure 1.**
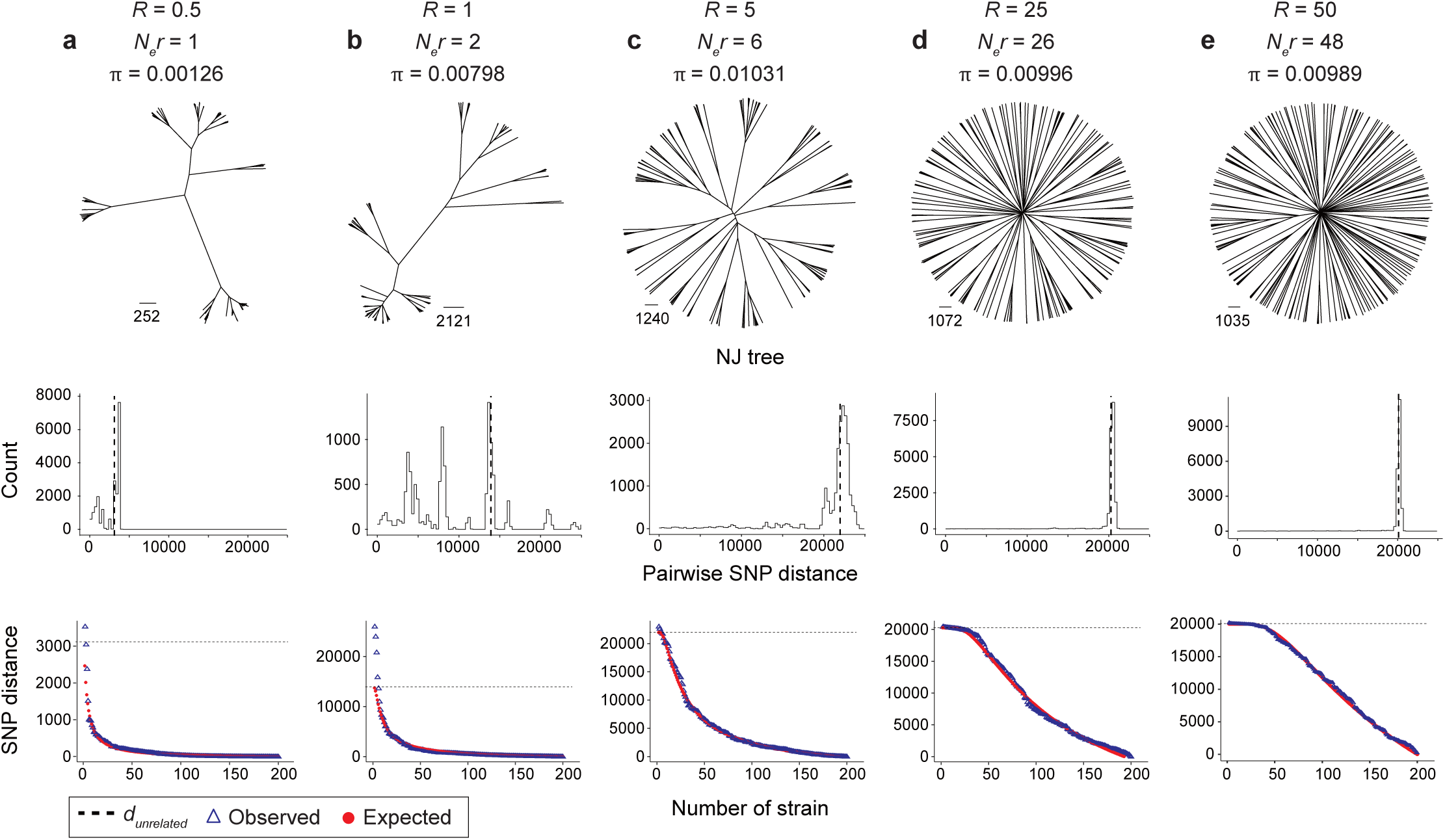
Recombination scaled effective population size (***N***_***e***_***r***) estimation of simulated constant populations under different scaled recombination rate. The panels indicated populations with scaled recombination rate *R* of 0.5 (a), 1 (b), 5 (c), 25 (d), 50 (e). From top to bottom, indicating the NJ trees, distribution of pairwise SNP distance between individuals and observed and expected coalescence curves. The dashed line of middle and bottom panels indicated the median SNP distance between individuals. The red points in the bottom panel indicated the expected distances between *n* - 1 coalescent nodes, the blue triangles indicated the observed distances estimated from pairwise SNP distances using the UPGMA algorithm.

Further simulations of more complex scenarios (Supplementary Fig. 1) show that *N*_*e*_*r* estimates reflect the current demography of the population more closely than estimates based on *π*, which is more influenced by past demography and migration to and from different demes. Supplementary Fig. 1a shows the effect of a population expansion on estimates of *N*_*e*_*r*. At time *t*, a single population with scaled recombination rate *R* = 5 splits into two. The blue population maintains the ancestral population size while the red population becomes 10 times larger. At time *t* + 0.05 the red population is only marginally more diverse than the blue one but its estimated value of *N*_*e*_*r* is 6 fold higher. The fit of observed and expected genetic distances is poor for the red population, reflecting the inaccuracy of the modelling assumption that there is a single unchanging population size. At time *t* + 0.2 the estimate of *N*_*e*_*r* is close to its true post-split value for both populations and the model fit is substantially improved. Merging data from the two populations gives intermediate estimates of *N*_*e*_*r*.

Supplementary Fig. 1b shows the effect of a population size reduction. A single population with *R* = 50 splits into two, with the blue population remaining unchanged while the red population undergoes a 10 fold reduction in size. As in Supplementary Fig. 1a, the model fit for the red population is poor immediately after the split but quickly improves, with the estimate of *N*_*e*_*r* approaching the correct value of 5, while the nucleotide diversity of the population reduces much more slowly. Supplementary Fig. 1c shows the effect of symmetric migration between two populations with a 10 fold difference in effective population size. Migration has a large effect on nucleotide diversity, especially for the smaller population, but has little effect on estimates of *N*_*e*_*r*. Overall, these results show that our inference approach provides accurate estimates of the recombination-scaled effective population size for simulated data and that deviations between observed and expected genetic distances can be informative about deviations from model assumptions.

### Application to ***V. parahaemolyticus* genomes**

We first applied the method to the 1,103 *V. parahaemolyticus* genomes described in Yang et al [17]. For this dataset, we obtained estimates of *N*_*e*_*r* = 484 and 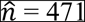 (Fig. 2a). 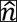 is less than half of the sample size principally because a large fraction of the isolates in the sample belong to pandemic clonal lineages responsible for large numbers of human infections. Both lineages have most recent common ancestors within the last few decades and are likely to represent a very small fraction of the global population of *V. parahaemolyticus*. The genetic distances fit the model well except that the 33 oldest coalescent events, on the left hand side of the plot are larger than *d*_unrelated_. The discrepancy is largely due to population structure within the species, since the fit is better, although still not perfect when analysis is restricted to the 944 isolates from the VppAsia population (Fig. 2a).

**Figure 2.**
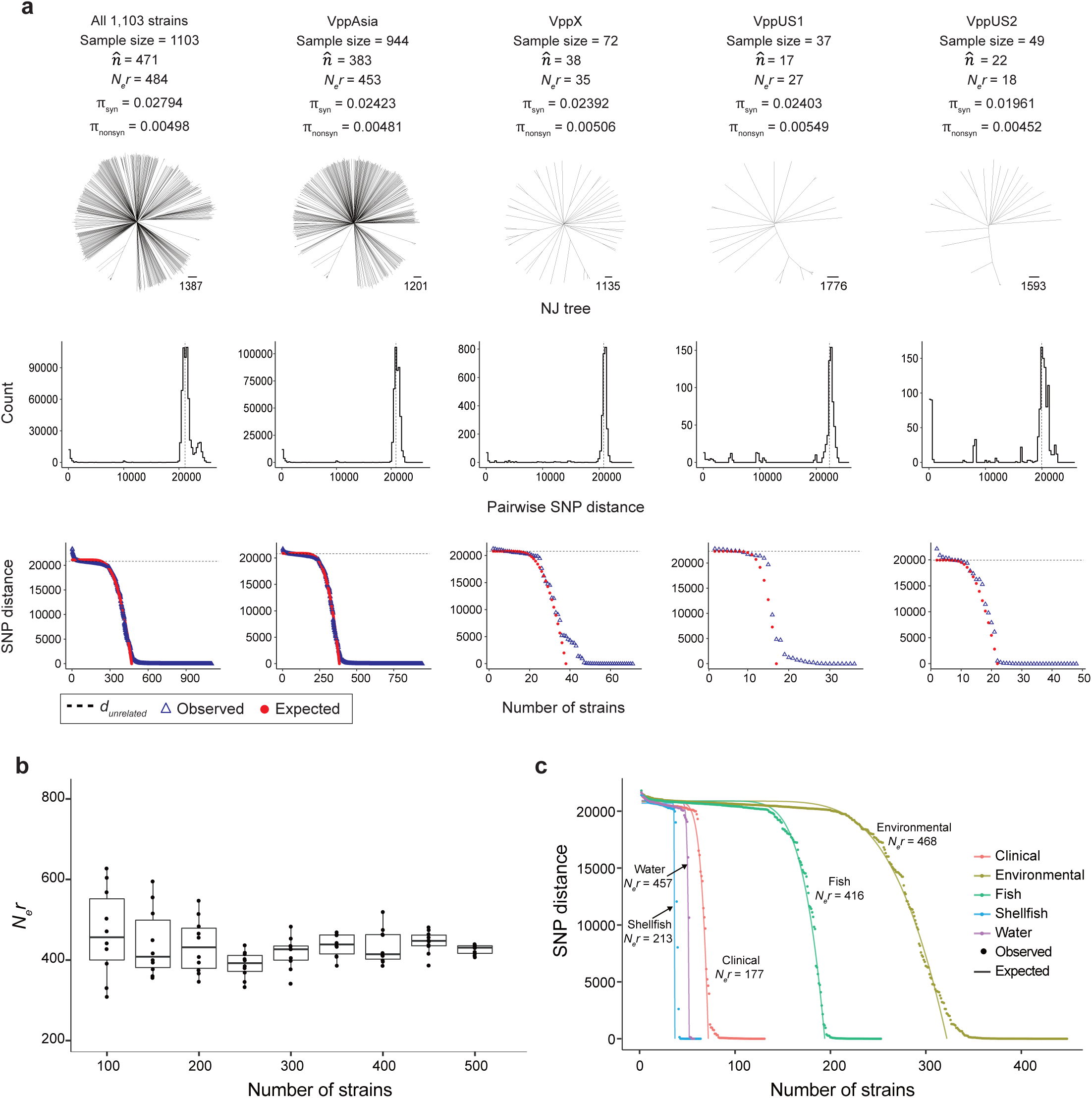
Recombination scaled effective population size (*N*_*e*_*r*) estimation of *V. parahaemolyticus*. (a) *N*_*e*_*r* estimation of all the samples and four populations (VppAsia, VppX, VppUS1 and VppUS2) of *V. parahaemolyticus*. Layout and colors are the same as in Figure 1. (b) *N*_*e*_*r* estimation of VppAsia population based on different sample sizes. 100-500 genomes were randomly selected from total 944 VppAsia genomes, 10 repeats were performed for each sample size to create the boxplot. (c) *N*_*e*_*r* estimation of VppAsia population based on different types of samples. Points and lines show observed and expected coalescence curves, respectively.

As described by Yang et al., the sample of *V. parahaemolyticus* is subdivided into 4 modestly differentiated populations, likely due to historical barriers to migration between oceans [17]. For the VppAsia population *N*_*e*_*r* was estimated to be 453 which is similar to that for the dataset as a whole, while other populations have substantially smaller values. These differences are not simply due to a larger sample size since estimates based on subsets of the VppAsia data are consistently greater than 200 (Fig. 2b). Sampling strategy does make some difference, since estimates are lower for a dataset consisting only of clinical strains than of shellfish, fish or all non-clinical isolates (Fig. 2c). This difference presumably reflects variation in disease causing potential, which results in samples of clinical isolates not representing the full diversity of clonal lineages within the species.

### Application to multiple bacterial species

We also applied the method to a survey of other bacteria for which large numbers of genomes are publically available (Supplementary Fig. 2). *V. cholerae* has an estimate of *N*_*e*_*r* of 29, while *Vibrio vulnificus* has a value of 43. Although the sample sizes available are relatively limited, in contrast to the Asian population of *V. parahaemolyticus*, the genetic distances do not show a clear plateau corresponding to a single value for *d*_unrelated_ and while there are a continuous range of coalescent distances, the overall fit between model and data is poor. Thus, although the other *Vibrio* species in our dataset recombine frequently they are far from being panmictic.

*H. pylori* has the largest estimated value of *N*_*e*_*r* of all datasets we analysed. *H. pylori* is characterized by extremely high rates of recombination during mixed infection of the human stomach, with 10% or more of the genome recombined during a single infection [23] and linkage disequilibrium decreases much more rapidly as a function of genetic distance than for all other species, including *V. parahaemolyticus* (Fig. 3a). Although the tree is approximately star-like, the fit of the model is not perfect, with no clear plateau for a single value of *d*_unrelated_, due to the complex geographical population structure of the species [24].

**Figure 3.**
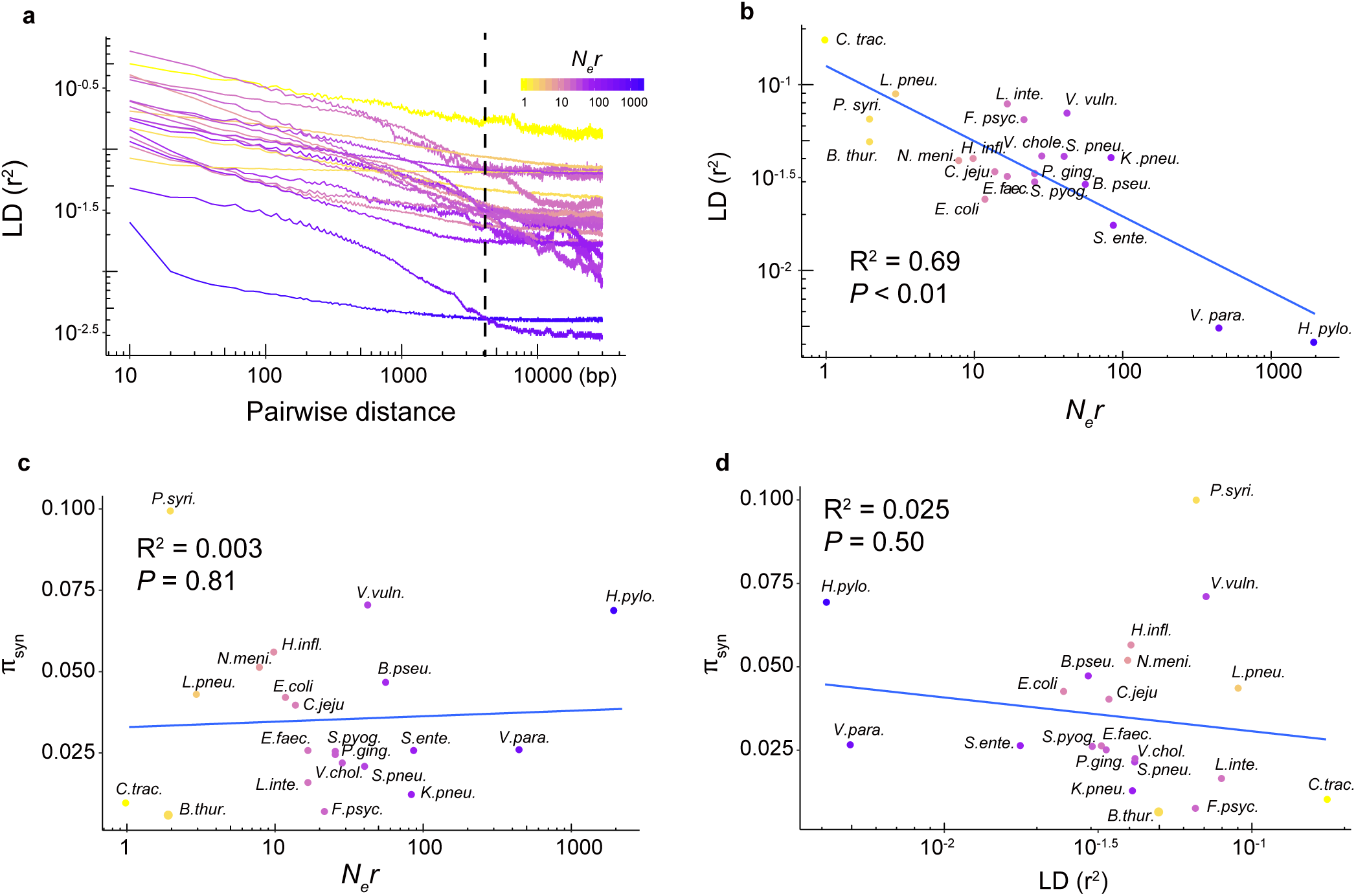
Correlation between *N*_*e*_*r*, linkage disequilibrium (LD) statistic *r*^2^ and nucleotide diversity of synonymous sites. (a) LD decay of 21 bacterial species. The maximum comparison distance was set to 30 kb. The vertical dashed line indicated the LD *r*^2^ values at pairwise distance of 3 kb, which were used in panel b and d. Line colours indicated the estimated *N*_*e*_*r* values. (b). Correlation between *N*_*e*_*r* and LD *r*^2^ values. (c) Correlation between *N*_*e*_*r* and nucleotide diversity of synonymous sites π_*syn*_. (d) Correlation between LD *r*^2^ values and nucleotide diversity of synonymous sites π_*syn*_. Point colours in panel b-d indicated the estimated *N*_*e*_*r* values.

Amongst the other species, *N*_*e*_*r* varies from 1 for *Chlamydia trachomatis* to 88 for *Salmonella enterica*. The overall fit of the model varies considerably between species (Supplementary Fig. 2) and is typically worst for the oldest coalescent events, as examined in more detail above for *V. parahaemolyticus*. Our estimated *N*_*e*_*r* values are correlated with the linkage disequilibrium statistic *r*^2^ measured between distant makers at pairwise distance of 3 kb (Fig. 3b). However, *N*_*e*_*r* shows no correlation with nucleotide diversity of synonymous sites *π*_*syn*_ (Fig. 3c) and *r*^2^ at pairwise distance of 3 kb also shows no correlation with *π*_*syn*_ (Fig. 3d).

## Discussion

### Structure of genetic diversity in 21 bacterial species

In bacteria, adaptation to diverse environmental challenges should be most effective in species where realized recombination rates are high enough to thoroughly mix up genetic variation (panmixia), since this creates the largest possible pool of genotypes on which natural selection can act. High recombination can also make it easier for researchers to detect the imprint of natural selection, a feature exploited by Cui et al. 2015, 2018 [8, 25] in investigating coadaptation in *Vibrio parahaemolyticus*. However, despite the utility of well-mixed gene pools, both to the species that have it and to the researchers studying it, it appears to be rare in bacteria for which large numbers of genomes are currently available.

The genetic structure of bacteria species depends on the interplay of many different processes, including changes in population size over time, geographical and ecological subdivision and the complex biology of genetic exchange which takes place via conjugation, transformation and transduction. As a result no single parameter summarizes the effect of recombination in breaking down linkage disequilibrium. Furthermore, available genomes rarely come close to being a random sample of the bacterial population from which they are taken.

Here we have used two summary statistics of the effectiveness of recombination; a non-parametric measure of long-range linkage disequilibrium, *r*^*2*^ between markers 3 kb apart on the genome, and a parametric approach estimating the composite population genetic parameter *N*_*e*_*r*. Informally, high values of *N*_*e*_*r* implies that recombination has generated new clonal complexes quickly compared to the rate at which genetic drift (proportional to 1/*N*_*e*_) removes them, with the result there are many distinct clonal-complexes segregating in the population. In data simulated according to a coalescent model with recombination of short tracts, estimated values of *N*_*e*_*r* are close to the true values for *N*_*e*_*r* > 2.

In our survey of 21 species from which large numbers of genomes are available, our two summary statistics are strongly but incompletely correlated (R^2^ = 0.69, *P* < 0.01, Fig. 3b). According to both statistics, two species are clear outliers, with estimates of *N*_*e*_*r* = 453 for the Asian population of *V. parahaemolyticus* and 1,976 for the hpEurope population of *H. pylori*, and the estimated value of *N*_*e*_*r* was lower than 100 in the other 19 species, which are therefore far from being panmictic.

Estimated *N*_*e*_*r* or *r*^*2*^ are uncorrelated with the nucleotide diversity of synonymous sites π_*syn*_ (Fig. 3c, Fig. 3d). For example, *V. parahaemolyticus* has similar diversity to *V. cholerae* and much lower diversity than *V. vulnificus*, both of which have estimated *N*_*e*_*r* lower than 50. Furthermore, estimates of *N*_*e*_*r* vary over a factor of 1,000, while *π* varies only by a factor of 10.

### Mechanisms by which high diversity can create barriers to recombination

Since panmixia is possible within bacterial populations and should facilitate genetic adaptation to the widest possible range of niches available to the species, it raises the question why it is not widespread. There are several bacterial species in our sample which survive well in the environment, have effective global dispersal, enormous census population sizes and high recombination rates, such as *E. coli, Campylobacter jejuni* and *K. pneumoniae*. Species with these characteristics are the most obvious candidates to be panmictic, whereas *V. parahaemolyticus* thrives only in warm brackish waters and has oceanic gene pools, implying historical limits on its dispersal [17], and *H. pylori* only survives in human stomachs and shows geographic differentiation associated with historical migrations of its host [26].

Neutral population genetic models imply that, all else being equal, bacteria with higher census population sizes should have higher *N*_*e*_*r* as well as higher nucleotide diversity, which is proportional to 2*N*_*e*_*μ* at equilibrium. However, across our 21 species, there is no correlation between the two statistics. We propose that the absence of a correlation occurs because high diversity tends to supresses recombination. There are a number of mechanisms by which this suppression can occur that have been described in the literature and we do not attempt to reach a conclusion about which are in fact most important in nature.

For example, in Cui et al [25], we observe nascent boundaries to genetic exchange between lineages of *V. parahaemolyticus* associated with differentiation into ecological types. This kind of genetic differentiation is more likely to be common, and more likely to lead to barriers to exchange in more diverse bacterial populations, which tend to have more diverse accessory genomes [27], and might also have a larger number of distinct ecological niches. Such barriers can result in speciation [21, 28] but can also lead to genetic structuring within a single species, such as host-specific gene pools found in *C. jejuni* [29].

Another mechanism is changes in the pattern or rate of recombination. Recombination requires homology between donor and recipient DNA to be recognized by the cellular machinery and, in addition, most species have a mismatch repair system, which aborts the process if there are too many differences. Therefore, as homology decreases, so will recombination [23, 28, 30, 31]. Mismatch dependent recombination is likely to be the main reason why *E. coli* has an estimated value of *N*_*e*_*r* of only 12, implying that recombination is ineffective in reassorting variation across the whole species [32].

Finally, a genetic dependence of recombination rates on diversity might arise from effects of epistasis in constraining realized recombination. Simulations of facultatively sexual organisms have shown that selection on multiple interacting loci can lead to populations being dominated by small numbers of clones, even in the presence of frequent recombination [33]. These simulations also show that a phase transition from panmixia can take place due to a decrease in recombination rate or even an increase in population size.

### *Vibrio parahaemolyticus* and *Helicobacter pylori* are far from demographic equilibrium

If, as we propose, suppression of recombination due to high diversity is a general phenomenon in bacteria, then the existence of populations with high *N*_*e*_*r* might seem paradoxical, because high *N*_*e*_ leads to high diversity which should suppress *r*. However this argument only holds in populations where *N*_*e*_ has been high for long enough for diversity to accumulate and do not apply if high *N*_*e*_ is a recent phenomenon on an evolutionary timescale.

In fact, neither the Asian population of *V. parahaemolyticus* nor *H. pylori* are close to demographic equilibrium. As proposed in Yang et al [17], the Asian population of *V. parahaemolyticus*, VppAsia, has spread within the last few decades due to human activity but has an ancestral range restricted to coastal waters from India to Japan. The other *V. parahaemolyticus* populations that we have sampled, with different ancestral ranges, have smaller estimated *N*_*e*_*r*, perhaps because they have smaller or less fecund ranges or experience greater competition or more frequent demographic bottlenecks.

Crucially, it seems that the ancestral population size of *V. parahaemolyticus* as a whole was smaller than currently found in VppAsia and more similar to that in the other populations. First, the site frequency spectrum of VppAsia, but not the other populations, is out of equilibrium and is approximately consistent with a demographic scenario in which the effective population size increased by a factor of ten 15,000 years ago (Supplementary Fig. 3). One possible scenario is that the end of the last ice age around 11,000 years ago created a habitat that existed till the present and that the population expansion implied by the site frequency spectrum is due to a demographic expansion at the end of the ice age.

Secondly the level of synonymous nucleotide diversity in the population is 30-fold lower than expected at demographic equilibrium. r/*μ*, the number of recombinant sites for each mutant site, has been estimated for the species to be around 313 [17]. Note that this parameter r/*μ* is not the same as the parameter r/*m* often used in the literature [6] because the former considers all recombined sites whereas the latter considers only recombinant sites that are substituted. If nucleotide diversity reached an equilibrium, consistent with the current value of *N*_*e*_*r* of VppAsia, this would predict a value of 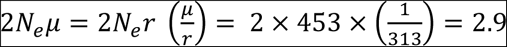. At equilibrium in a neutrally evolving population, with sites evolving according to the Jukes-Cantor model, *π* is predicted to be equal to 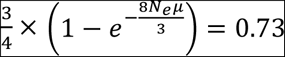, and therefore most synonymous sites should differ between individuals. This is far from the current value of 0.024 or indeed diversity levels in any well-characterised bacterial species. Our simulation results show that estimates of *N*_*e*_*r* respond much more quickly to changes in demography than *π*, implying that diversity levels would be likely to continue a slow but steady rise if the effective population size remains constant (ignoring the changes in population structure caused by human activity in recent decades).

The equivalent argument is more complex in *H. pylori* due to its geographic population structure but levels of variation are clearly far from demographic equilibrium, as illustrated by the substantial variation between populations that reflects bottlenecks associated with historical human migration rather than current population sizes [26]. These results therefore suggest that if effective population sizes remain continuously high in either species, then diversity would increase. We propose that in this case, barriers to recombination would evolve, perhaps by one of the mechanisms described above.

Despite decades of study by population geneticists, the factors determining the amount of diversity within species are still poorly understood, for example with much lower variation in nucleotide variation *π* between species than is expected based on variation in census population size in many domains of life [34, 35]. Our results suggest that patterns of genome variation in bacteria are likely to be dynamic, with ecological and genomic differentiation, barriers to gene flow and demographic factors interacting with each other in complex ways to alter both the level of genetic diversity within the species and the way it is partitioned, to produce the wide variety of bacterial population structures that are observed.

## Material and methods

### Genomic datasets used in analysis

Because our inference approach of *N*_*e*_*r* hypothesizes that recombination drives the genetic variation other than mutation, we firstly selected 27 bacterial species, in which r/*m* values (ratio of nucleotide changes resulted from recombination relative to point mutation) of them were greater than one based on previous multi-locus sequence typing data estimation [6]. We then counted the number of assembled genomes of them in NCBI database, and found that the genome numbers of 21 bacterial species were greater than 100, which were used in further analysis, including *Bacillus thuringiensis, Burkholderia pseudomallei, Campylobacter jejuni, Chlamydia trachomatis, Enterococcus faecalis, Escherichia coli, Flavobacterium psychrophilum, Haemophilus influenzae, Helicobacter pylori, Klebsiella pneumoniae, Legionella pneumophila, Leptospira interrogans, Neisseria meningitidis, Porphyromonas gingivalis, Pseudomonas syringae, Salmonella enterica, Streptococcus pneumoniae, Streptococcus pyogenes, Vibrio cholerae, Vibrio parahaemolyticus* and *Vibrio vulnificus.*

For species with more than 500 genomes in NCBI database, such as *E. coli* (>14,000 assembled genomes), 500 genomes were randomly selected to reduce the amount of calculation. Each genome was aligned against the reference genome of the corresponding species (Supplementary Table 1) using MUMmer [36] to generate the whole genome alignments and identify SNPs in core genomes (regions presented in all isolates) as previously described [8, 17]. Only genomes with genome-wide coverage > 70% (compared to reference genome) were used in further analysis. SNPs located in repetitive regions were removed, and the filtered bi-allelic SNPs sets were used to construct the Neighbour-joining (NJ) tree of each species. Strains located on the extremely long branches of the NJ tree and strains belonged to clonal groups were manually removed, finally resulting in a dataset of 6,355 genomes (56-1,103 genomes for each species). The accession numbers of genomes used (excluding *H. pylori* and *V. parahaemolyticus*) were listed in Supplementary Table 1, and the whole-genome alignments were available in the figshare data repository (https://figshare.com/s/3f9d04a8229f30dd785b).

### SNP calling, phylogeny reconstruction and LD decay calculation

The SNP dataset of 278 *H. pylori* (hpEurope population) and 1,103 *V. parahaemolyticus* genomes were reused from previous studies [17, 37]. The SNPs of the left 4,974 genomes were recalled using same pipelines as described above. Totally 17,875-462,214 bi-allelic SNPs were identified separately for 21 bacterial species, which were used in further analysis, including Neighbour-joining tree construction, pairwise SNP distance calculation, LD *r*^*2*^ value calculation, and *N*_*e*_*r* estimation as previously described [8]. The Neighbour-joining trees were constructed using the software Treebest (http://treesoft.sourceforge.net/treebest.shtml) based on concatenated SNPs. Haploview [38] was used to calculate the LD *r*^*2*^, and the maximum comparison distance was set to 30 kb.

## Simulation

The software FastSimBac [22] was used to generate the simulated bacterial populations under different hypothetical evolution scenarios, including a constant population with different scaled recombination rate *R* (Fig. 1), and the changing populations with population expansion (Supplementary Fig. 1a), reduction (Supplementary Fig. 1b) and migration (Supplementary Fig. 1c). All the simulated genome length was set to 2 Mb, mutation rate was fixed at 0.01 (per site per 2*N*_*e*_ generations). The detailed parameters used in simulation were listed in Supplementary Table 2.

### Recombination scaled effective population size estimation

For a freely mixing population in which recombination drives diversification, neutral theory predicts that *N*_*e*_*r* is in proportion to genealogical coalescent rate, and the expected coalescent curves can be estimated based on the formula 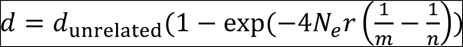 as mentioned in previous work [8]. In the formula, *d* is the expected pairwise genetic divergence, *d*_unrelated_ is the median pairwise SNP distance of a population, *n* is the number of individual strains and *m* is the index of the ancestral node along the coalescent tree of *n* strains. Coalescent curves can be estimated using the UPGMA algorithm based on the pairwise SNP distance. By fitting the expected and observed coalescent curves with least square method to search for the optimal parameters, we found the optimal values of 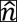 (effective sample size) and *N*_*e*_*r* that were used in further analysis.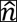 was an estimate of the number of strains remaining when over-sampled clonally related strains are removed.

### Site frequency spectrum estimation

The minor allele frequency (MAF) of each SNP locus in a SNP matrix was calculated and then the frequency of SNP positions at each MAF level were counted to generate the site frequency spectrum (SFS) for each dataset. To get a comparable result, the SFS showed in each Figure or panel was calculated based on same sample size. In Supplementary Fig. 3a, the merged population was generated by randomly selecting 100 genomes from each of Pop1 and Pop2. In Supplementary Fig. 3b, the same number of genomes as in the real genome dataset were simulated to show populations with different evolution scenarios.

## Acknowledgements

This work is supported by the National Key Research & Development Program of China (No. 2017YFC1200800, 2017YFC1601503, and 2016YFC1200100), the National Key Program for Infectious Diseases of China (No. 2017ZX10104002), the National Natural Science Foundation of China (No. 31770001) and Sanming Project of Medicine in Shenzhen (No. SZSM201811071). D.F. is funded by a Medical Research Council Fellowship as part of the MRC CLIMB consortium for microbial bioinformatics (grant number MR/M501608/1).

## Author Contributions

D. F., Y. C., and R. Y. designed the study and coordinated the project; C. Y., Y. C., X. D., and D. F. analyzed the data; D. F. wrote the manuscript. All authors approved the final version of the manuscript.

