## Supplementary Figures for "Why panmictic bacteria are rare"

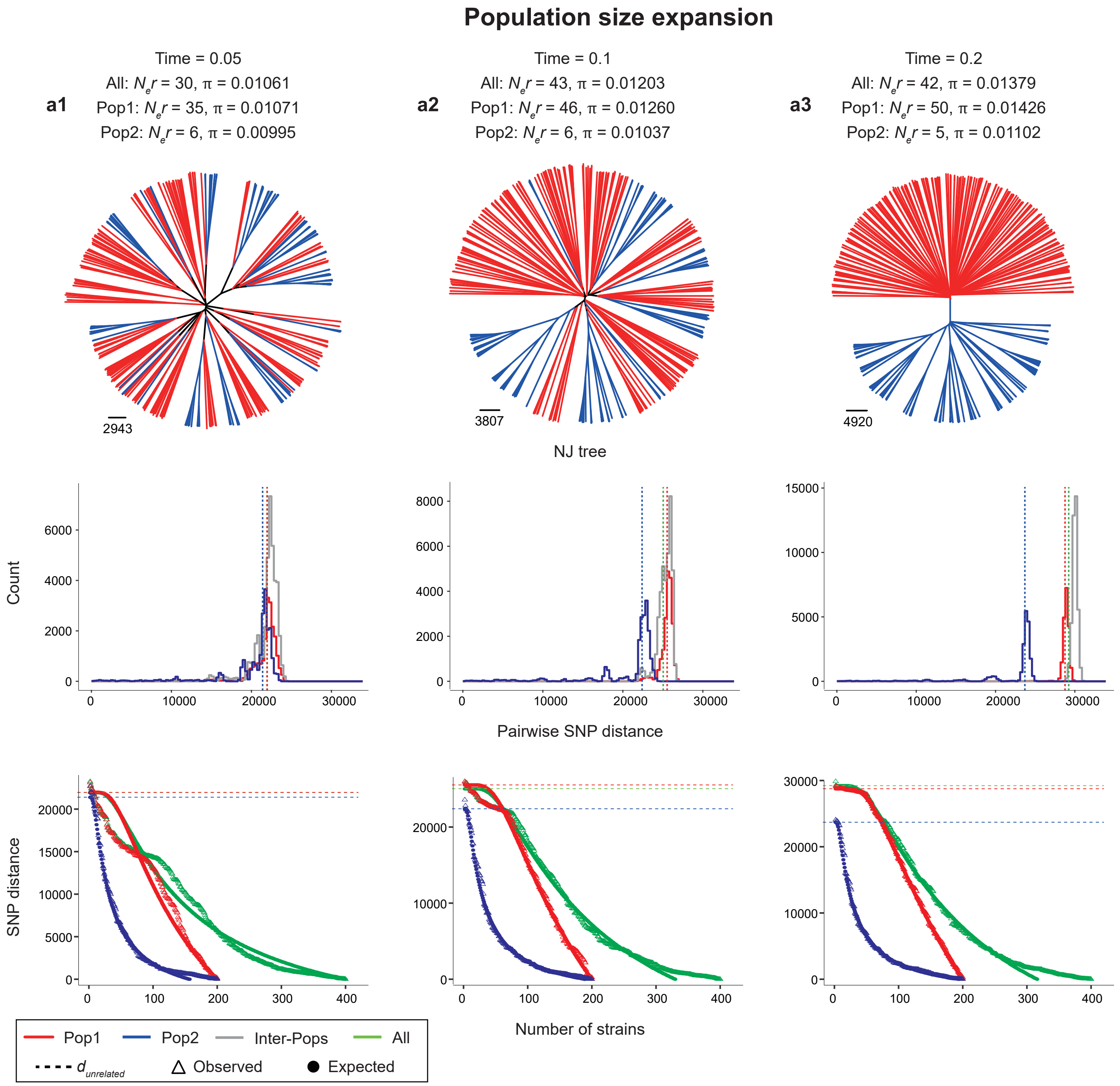


**Supplementary Figure 1. Recombination scaled effective population size ($N_er$) estimation of simulated populations with different dynamics patterns: population size expansion (a), reduction (b) and migration (c).** Panel a indicated population size expanded 10-fold at the time of 0.05 (a1), 0.1 (a2) and 0.2 (a3). Panel b indicated population shrank 10-fold with different timescales of 0.001 (b1), 0.01 (b2) and 0.05 (b3). Panel c indicated population migrated at the different rates with scale of 1 (c1), 5 (c2) and 10 (c3) at the time of 1. From top to bottom, indicating the NJ trees, distribution of pairwise SNP distance between individuals and observed and expected coalescence curves. The dashed line of middle and bottom panels indicated the median SNP distance between individuals. The red points in the bottom panel indicated the expected distances between n-1 coalescent nodes, the blue triangles indicated the observed distances estimated from pairwise SNP distances using the UPGMA algorithm. Pop1 indicates the population with changed $N_e$ and were marked in red. Pop2 indicates the original population with stable $N_e$ and were marked in blue. Inter-populations distances and $N_er$ estimation of all samples were marked in grey and green, respectively.

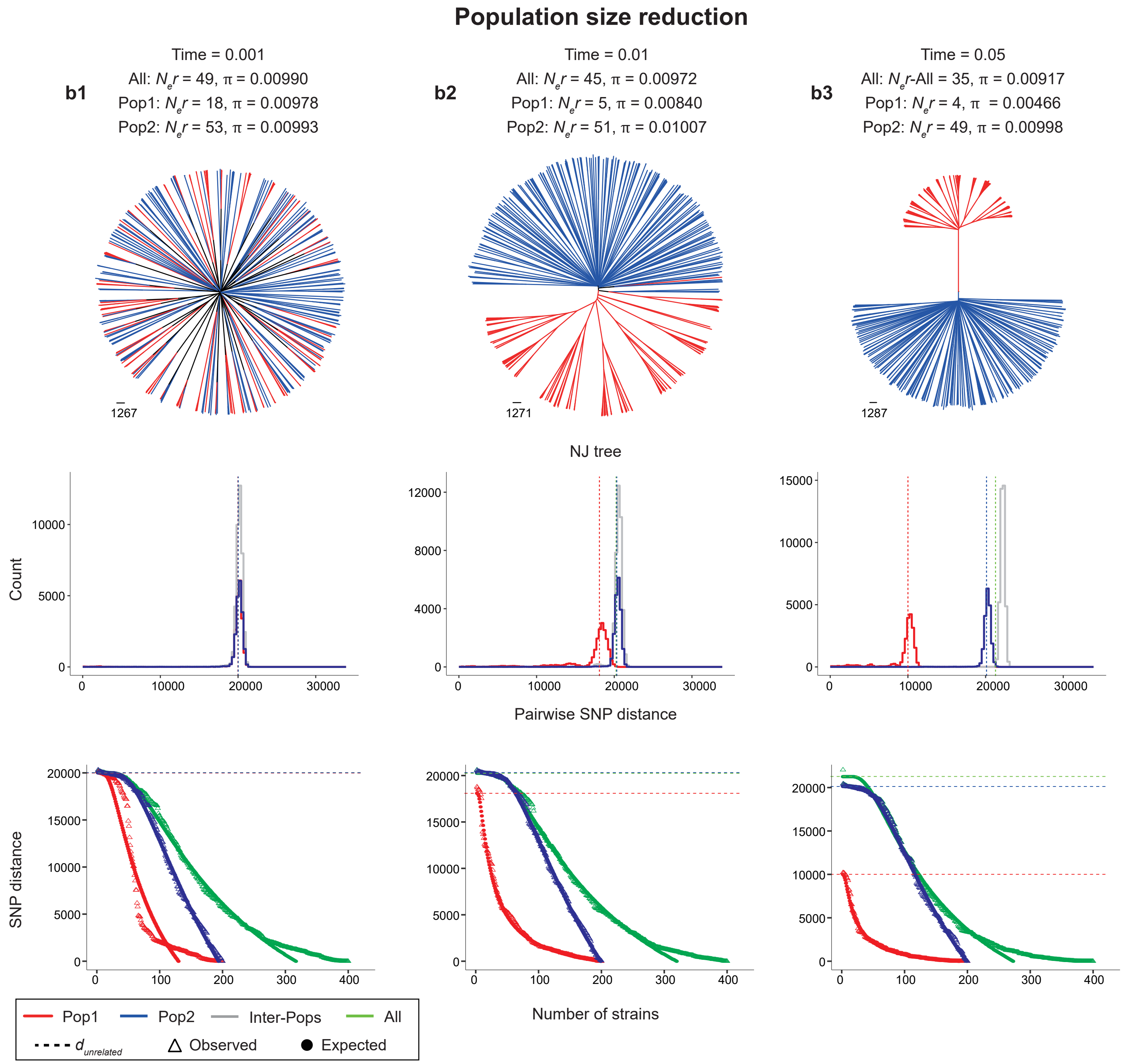


**Supplementary Figrue 1 continued.**

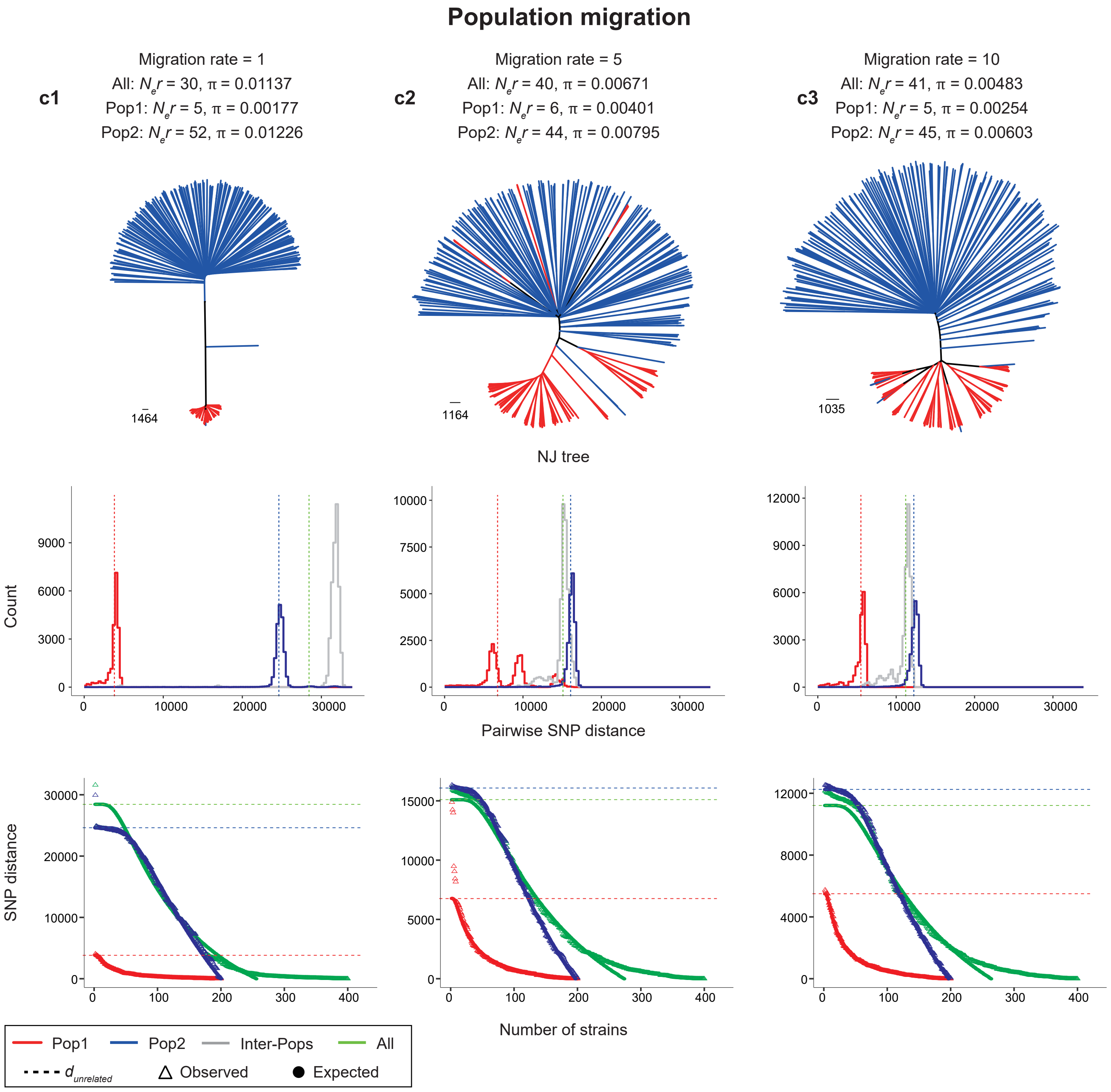


**Supplementary Figrue 1 continued.**

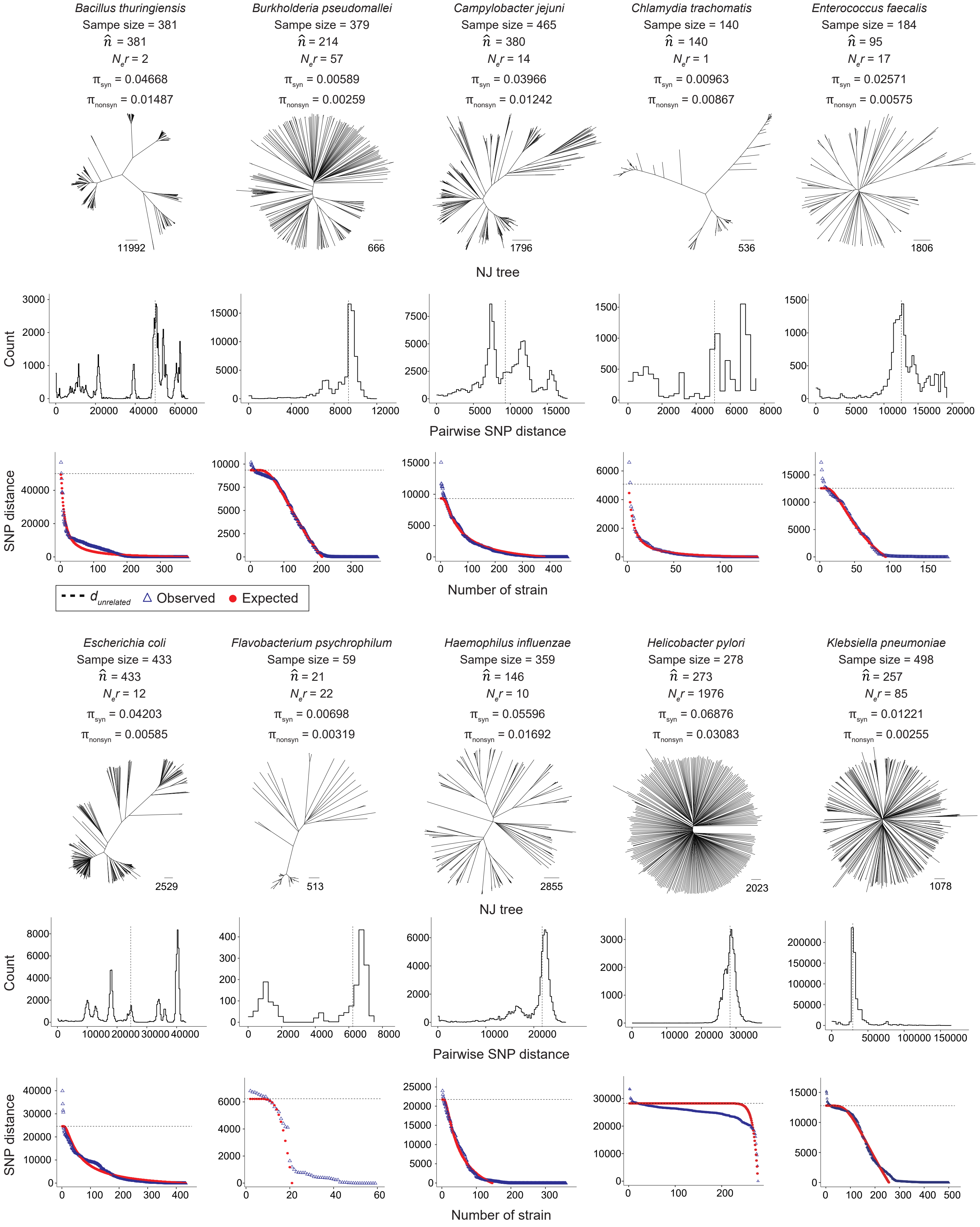


**Supplementary Figure 2. Recombination scaled effective population size ($N_e r$) estimation of 20 other bacterial species.** Layout and colors are the same as in Figure 1 and Figure 2a. From top to bottom, indicating the NJ trees, distribution of pairwise SNP distance between individuals and observed and expected coalescence curves. The dashed line of middle and bottom panels indicated the median SNP distance between individuals. The red points in the bottom panel indicated the expected distances between n-1 coalescent nodes, the blue triangles indicated the observed distances estimated from pairwise SNP distances using the UPGMA algorithm.

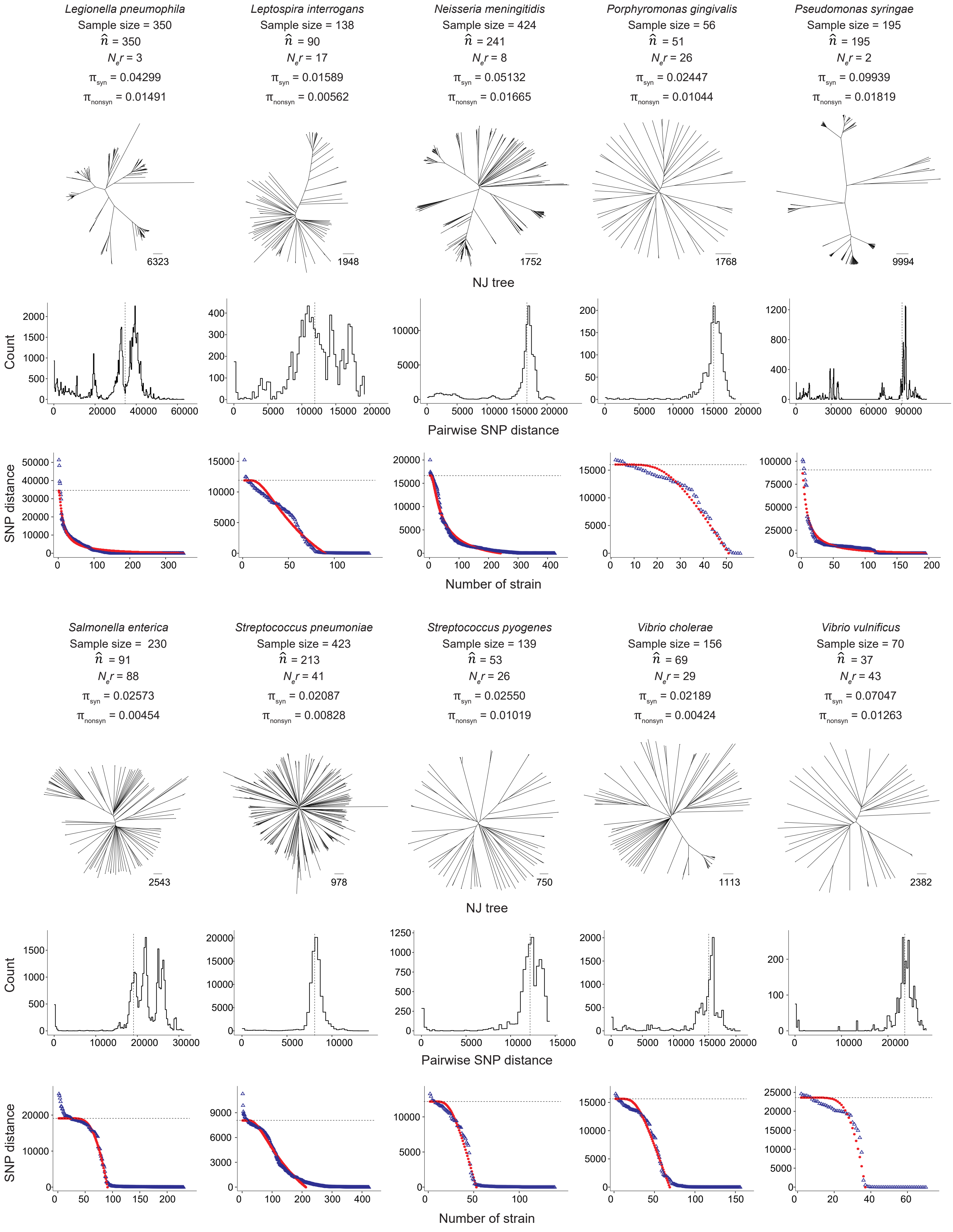


**Supplementary Figure 2 continued**

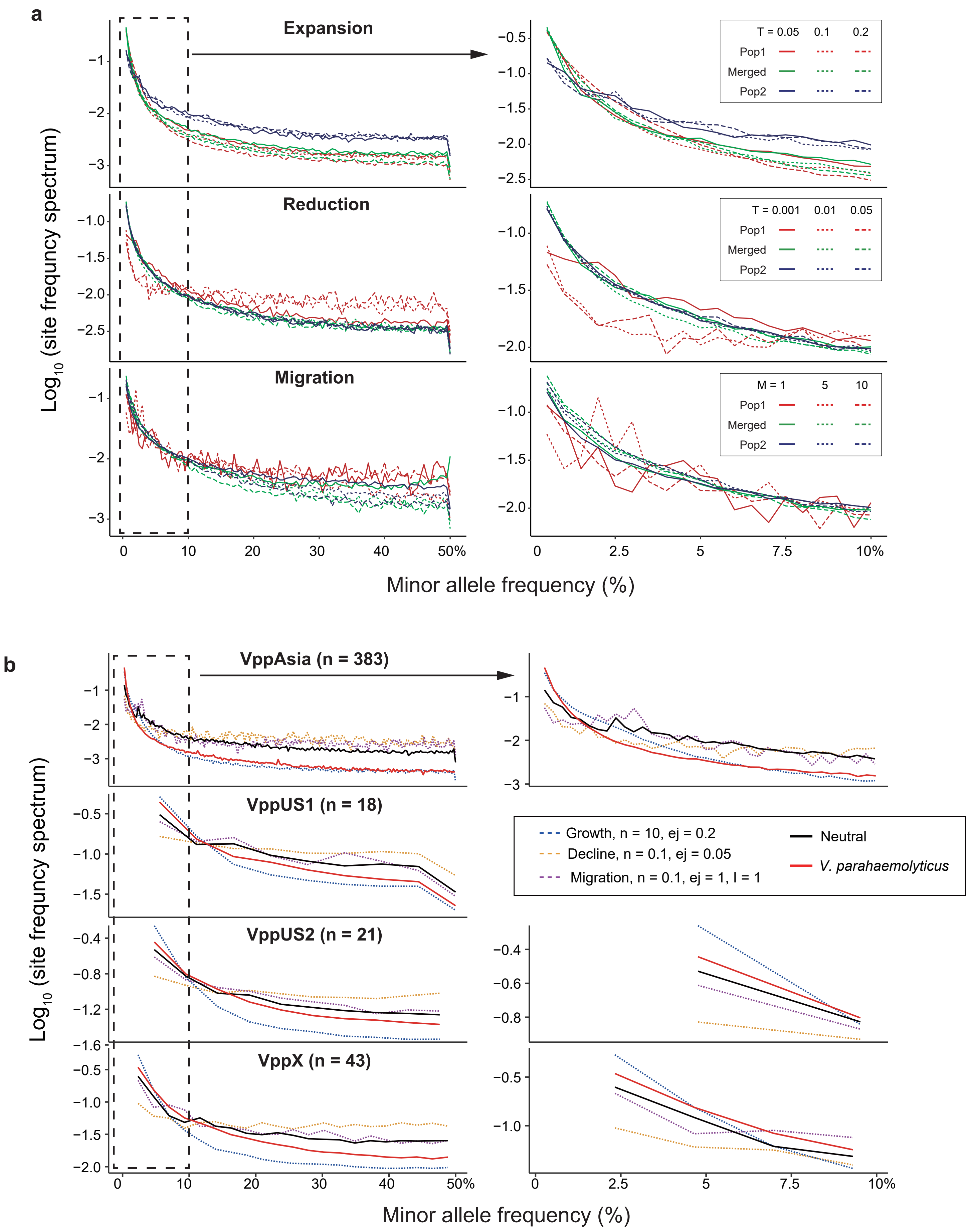


**Supplementary Figure 3. Site frequency spectrum of simulated populations and *V. parahaemolyticus* populations.** (a) From top to bottom, indicating simulated population size expansion, reduction and migration, corresponding to Supplementary Figure 1a-1c. Pop1 indicates the changed population and were marked in red. Pop2 indicates the original populations with stable $N_e$ and were marked in blue. Merged population (generated by randomly selecting 100 genomes from Pop1 and Pop2, respectively) was marked in green. Line types show the time of population size change. (b) Site frequency spectrum of four *V. parahaemolyticus* populations. Only non-redundant strains (no sequences differed by less than 2,000 SNPs) were used to minimize the influence of clonal groups. For each *V. parahaemolyticus* population, same number of genomes were simulated to represent populations of different dynamics patterns. The right hand panel was the zoomed view of the minor allele frequency <10%.
